# Analysis of haemolysis-based injury in immortalised proximal tubule epithelial cells is impaired by haemoglobin interference with bioluminescence assays

**DOI:** 10.64898/2026.11.06.764891

**Authors:** Charlotte Trott, James Bennett, Patricia A. Murray, Amy Chadwick, Stephen J. McWilliam, Bettina Wilm

## Abstract

Acute kidney injury (AKI) is a serious complication which is common amongst cardiac and aortic/vascular surgery patients. Although the cause of AKI following cardiac surgery is multifactorial, haemolysis is an increasingly recognised contributing factor caused by the use of cardiopulmonary bypass machines. Haemolysis, the lysis of red blood cells, releases free haemoglobin (fHb) into the bloodstream which is known to cause kidney injury. Currently, the mechanisms of haemolysis-induced kidney injury are not fully understood. In this study, we investigated the injury response to fHb in a human-derived conditionally immortalised proximal tubule epithelial cell line (ciPTECs). Results showed that fHb interference reduces bioluminescence signal output which is considerably mitigated by removing fHb solutions and haemolysed plasma prior to the beginning of the assay. Interestingly, we found that the effect of fHb on ciPTECs was different with haemolysed plasma compared to fHb solutions, potentially due to factors other than fHb within the plasma. In contrast to expectations, haemolysed plasma induced only a marginal increase in NF-ĸB activity and had minimal effect on mitochondrial function. Together, these findings suggest that caution should be applied when using bioluminescent assays with fHb, both in solution and haemolysed plasma, and that fHb may not inflict severe injury on ciPTECs.

**New & Noteworthy:** fHb impairs the use of bioluminescent assays when investigating haemolysis-induced acute kidney injury. Removal of fHb restores bioluminescent signal output, revealing that fHb has minimal effect on a proximal tubule epithelial cell model.

## Introduction

Acute kidney injury (AKI) is common in patients following cardiac and aortic surgery requiring cardiopulmonary bypass (CPB), with a reported incidence between 20 – 30 % [1, 2]. AKI is a sudden, significant reduction in kidney function, and is associated with increased length of patient stay in hospital, mortality risk and the likelihood of requiring renal replacement therapy (RRT) [2, 3], as well as increasing the long-term risk of developing chronic kidney disease [4]. The cause of CPB-induced AKI is multifactorial, including renal hypoperfusion, hypoxia, inflammation and oxidative stress [5]. Haemolysis is considered another contributing factor; the lysis of red blood cells occurs due to the mechanical shear stress during the use of CPB, leading to the release of fHb [6]. Previous studies have shown that higher plasma fHb levels following cardiac surgery are associated with AKI development [7-9].

During cardiac and aortic surgery, the haemolysis-induced release of fHb can overwhelm the body’s natural scavenging system consisting of haptoglobin and hemopexin, increasing circulating levels of fHb and its breakdown products free haem and iron [10, 11]. These are primarily cleared from the bloodstream via glomerular filtration and taken up by the proximal tubule epithelial cells where they disrupt their function, leading to kidney injury [10, 12]. Identified mechanisms of injury include lipid peroxidation, oxidative stress, mitochondrial dysfunction and the induction of inflammation and cell death pathways [10, 12].

Here we analysed the effect of fHb in a proximal tubule cell model with the aim to investigate mechanisms of haemolysis-induced injury. We exposed a human-derived conditionally immortalised kidney proximal tubule epithelial cell line, ciPTECs, to a range of fHb concentrations similar to those observed in the blood of cardiac surgery patients following CPB, as well as haemolysed plasma from healthy donors. Our results indicate experimental challenges when investigating the biological effects of fHb in an in vitro model of haemolysis-induced AKI since Hb strongly absorbs light across the visible spectrum [13, 14], potentially interfering with commonly used laboratory assays.

## Materials and methods

### Samples

Blood samples were collected from aortic surgery patients pre-operatively, upon entry into surgical theatre before CPB was commenced, and post-operatively following weaning from CPB and administration of protamine sulphate. These samples were obtained under the HaemoAKI project which was approved by the Hampstead Research Ethics Committee (22/PR/0933).

Whole blood was collected from healthy volunteers under the Pharmacology Biobank project which was approved by Liverpool University’s Central University Research Ethics Committee D (11499). All participants gave written consent in line with Good Clinical Practice and declaration of Helsinki.

### Cell culture

ciPTECs (Cell4Pharma, Oss, NL) [15] were cultured in Dulbecco’s modified eagle’s medium and nutrient mixture F-12, phenol red free (DMEM/F12, Gibco,11039021) supplemented with Insulin (5µg/ml) -transferrin (5µg/ml) -sodium selenium (5ng/ml) (Sigma-Aldrich, I1884), 36ng/ml hydrocortisone (Sigma-Aldrich, H0135), 10ng/ml Epidermal Growth Factor (EGF, Sigma-Aldrich, 9644), 40pg/ml tri-iodothyronine (Sigma-Aldrich, T5516), 10% FBS (Gibco, A5256701). Cells were maintained at 33°C, 5% CO_2_. Medium was changed every 2 – 3 days.

Cells were passaged upon reaching 80-90% confluency. For experiments, cells were seeded at a density of 55,000 cells/cm^2^ in 96-well plates, unless otherwise stated. Following seeding, cells were cultured at 33°C, 5% CO_2_ for 24 hours (h) before being transferred to 37°C, 5% CO_2_ for 7 days to allow for differentiation. Medium was changed every 2 – 3 days. Cells were used between passages 35 and 55.

### Manual haemolysis of healthy volunteer plasma

Half of the healthy volunteer blood was haemolysed by passing it through a 16G needle multiple times. The remaining half remained un-haemolysed to serve as baseline plasma. Both blood preparations were centrifuged for 10 min at 3,000 g (Supplementary Figure S1). Plasma was carefully removed, aliquoted and stored at -80°C.

### Viability assessment

A 40 mg/ml stock solution of human Haemoglobin A_0_ (Sigma-Aldrich, H0267) (called fHb-A_0_ from hereon) in cell culture medium was filtered through a 0.22 µm PES membrane filter, aliquoted and frozen at -20 °C. ciPTECs were treated with fHb-A_0_ concentrations at 0.1 to 10 µg/µl or human plasma for 24 h or 48 h. In some experiments, treatment media or plasma was replaced with fresh medium immediately prior to the assay. CellTiter-Glo (Promega G7570) or CellTiter-Glo 2.0 (Promega, G9242) reagent was added in a 2:1 ratio of medium to reagent. The plate was shaken for 2 min followed by 10 min incubation at room temperature in the dark. Assay solutions were transferred to white 96-well plates and bioluminescent signal measured by BMG LABTECH FLUOstar omega plate reader (Supplementary Figure S2).

### Light Microscopy

Light microscopy imaging of ciPTECs treated with aortic surgery patient plasma was performed on the Leica DM500 microscope using the Leica DFC420 C camera, and images adjusted in ImageJ [16].

### Generation of ciPTEC NF-ĸB reporter line

A stable NF-κB reporter ciPTEC line was generated by transduction with lentivirus obtained using the pGreenFire1-NF-κB reporter construct, envelope plasmid (pMD2.G) and packaging plasmid (psPAX2), following standard protocols [17]. Cells were treated overnight with a solution of 8µg/ml polybrene and NF-ĸB Puromycin lentiviral particles, to give a Multiplicity of Infection of 10. Media was replaced the following day. After 2 days, transduced cells were selected with 0.5 μg/ml puromycin (InvivoGen, ant-pr-1), determined by optimisation of puromycin selection conditions, for 10 days and expanded. Reporter expression was confirmed by GFP fluorescence measured by flow cytometry (BD FACScalibur), while reporter functionality was validated by TNF-α stimulation (100 ng/ml, 24 h), which resulted in increased luciferase activity after addition of D-luciferin (Promega, E1605) (BMG LABTECH FLUOstar omega plate reader).

### NF-ĸB activity assessment

The ciPTEC-NF-ĸB reporter line was treated with human plasma or 100 ng/ml TNF-α (PeproTech, 30001A) for 6 or 24 h. Treatment was removed and replaced with fresh medium immediately prior to the assay. D-luciferin was added with a final concentration 200 µg/ml and luciferase expression quantified using the BMG LABTECH FLUOstar omega plate reader and analysed using Flo.reada. Background expression was subtracted, expression normalised to the untreated control and expressed as fold-change.

### Mitochondrial stress assessment

ciPTECs were seeded at a density of 10,000 cells/well in Seahorse XFe 96-well microplates (Agilent, 103794-100). Following differentiation ciPTECs were treated with human plasma for 24 h. Prior to the assay, plasma was removed and cells washed with seahorse complete medium, seahorse XF DMEM (Agilent, 103680-100) supplemented with 2 mM L-glutamine, 25 mM D-Glucose and 1 mM sodium pyruvate. 175 µl of seahorse complete medium was added and cells were incubated at 37 °C in the absence of CO_2_ for 1 h.

Oxygen consumption rate (OCR) and extracellular acidification rate (ECAR) were measured in realtime using the Seahorse XFe instrument (Agilent) using the Mito Stress Test protocol. Briefly, following the initial three basal measurement cycles, mito stress test compounds oligomycin (1.25 µM), carbonyl cyanide p-trifluoromethoxy phenylhydrazone (FCCP) (0.5 µM), rotenone and antimycin (1 µM each) were sequentially added. Three measurement cycles were performed after each addition.

BCA assay was used to normalise the OCR (pmol/min) and (ECAR (mpH/min) to protein concentrations. These values were then used to calculate parameters of mitochondrial function (Supplementary Table S1).

### Statistical Analysis

All statistical analysis was performed on GraphPad Prism 10 software and all data were normalised to the media control and presented as mean ± standard deviation (SD). Paired, two-tailed t-tests (two groups) and One-way ANOVA followed by Dunnett’s (all groups compared to a control group) or Tukey’s (all groups compared to each other) multiple comparisons were used. P<0.05 was considered statistically significant.

## Results

To investigate the effect of haemolysis on ciPTECs, cell viability was measured in response to 24 h treatment with fHb-A_0_ solutions or haemolysed plasma. Our initial dose-finding experiment using a range of fHb-A_0_ concentrations revealed a concentration-dependent reduction in ciPTEC viability, with an IC50 of 0.5 µg/µl fHb-A_0_ (Figure 1A). When ciPTECs were treated with manually haemolysed plasma from 4 healthy volunteers, cell viability was significantly reduced by approximately 50 % compared to the respective baseline plasma. Mean ciPTEC viability was 93.1 % ± 4.5 when treated with baseline plasma, while treatment with haemolysed plasma reduced viability to 49.4 % ± 7.3 (Figure 1B). The concentration of fHb within haemolysed plasma samples was measured using the Hemcheck device which demonstrated that the extent of ciPTEC viability loss generally correlated with fHb concentrations within the plasma (Figure 1C & Supplementary Table S2).

**Figure 1.**
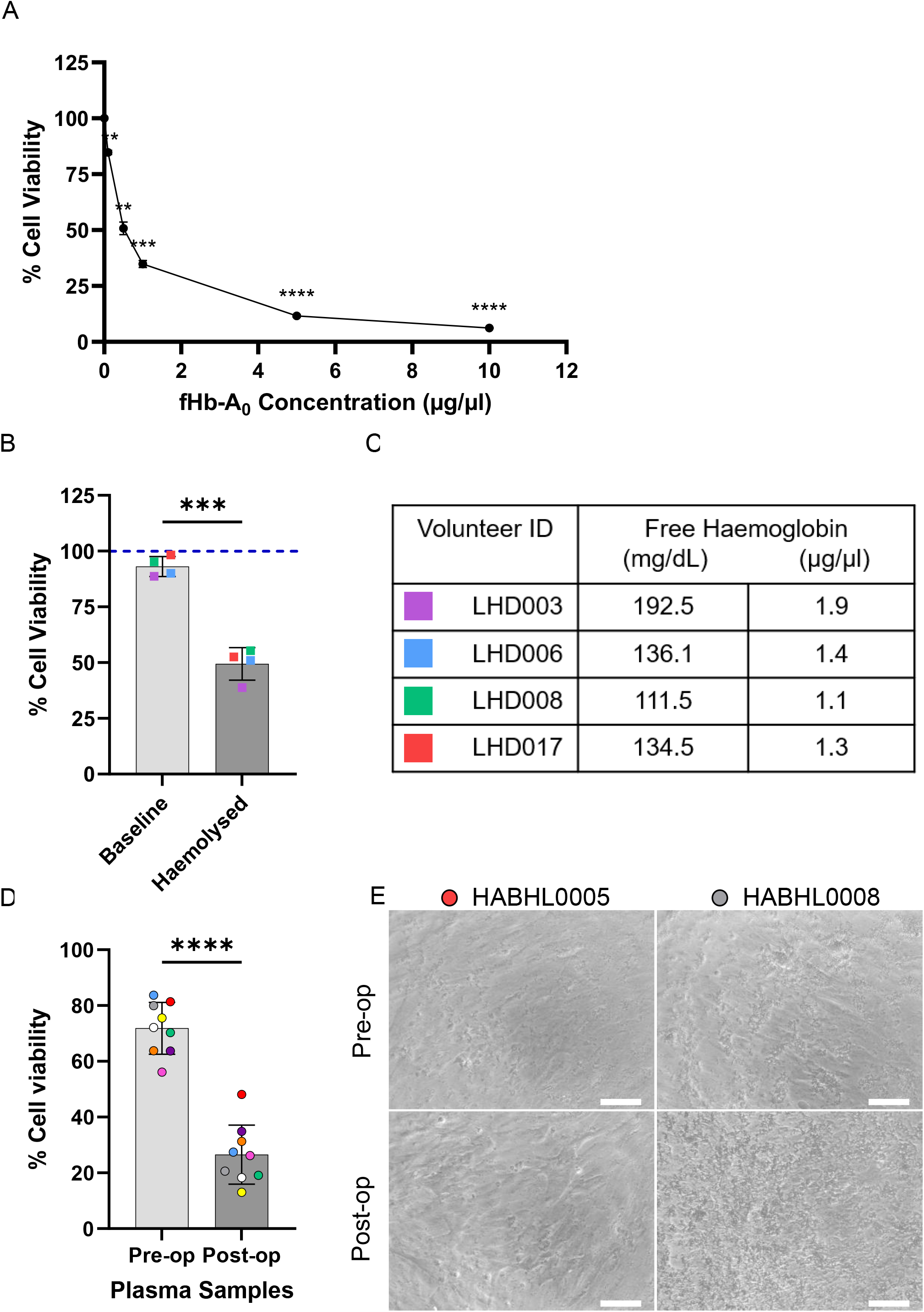
Investigating the effect of haemolysis on ciPTEC viability. ciPTECs were treated for 24 h with (A) a range of free haemoglobin (fHb-A_0_) concentrations from 0.1 to 10 µg/µl, (B) baseline and manually haemolysed plasma from 4 healthy volunteers and (D) pre- and post-operative plasma from 9 aortic surgery patients. The CellTiter-Glo assay was used to determine cell viability based on cellular ATP content. Data presented is from 3 independent experiments shown as mean % ± SD of the media control. In (B) media control is represented by the dotted blue line and individual data points represent individual volunteers, in (D) individual data points represent individual patients and all are colour-coded. Statistical significance in (A) from the media control was determined by One-Way repeated measures ANOVA with Dunnett’s multiple comparisons test and is represented by *, p<0.05, **, p<0.01, ***, p<0.001, ****, p<0.0001. Statistical significance in (B & D) was determined by paired t-test and is represented by ***, p<0.001, ****, p<0.0001. (C) fHb concentrations measured, by Hemcheck, of the haemolysed plasma samples used in (B). (E) Representative images of ciPTECs following 24 h treatment with patient plasma used in (D), scale bars represent 100 µm.

Next, we tested pre- (pre-op) and post-operative (post-op) plasma samples from 9 adult aortic surgery patients, where massive haemolysis can occur [18]. Treating ciPTECs with patient pre- and post-op plasma produced similar results to volunteer baseline and haemolysed plasma. Mean cell viability was significantly reduced by more than 50% from 71.9% ± 9.3 (pre-op plasma) to 26.6% ± 10.6 (post-op plasma; Figure 1D), suggesting considerable loss by cell death. Light microscopy analysis of ciPTECs treated with samples from all aortic surgery patients, revealed that varying quantities of visible debris were present in both pre- and post-op plasma (Figure 1E). However, post-op plasma with higher fHb concentrations (i.e. from patient HABHL0008), showed high quantities of debris which prevented observation of the cells. Where cells remained visible after treatment with post-op plasma, such as patient HABHL0005, we did not observe any change in ciPTEC morphology between pre- and post-op plasma treatment (Figure 1E).

The lack of changes in ciPTEC morphology despite such drastic reductions in cell viability after treatment with post-op plasma samples suggested that the viability results may not have been accurate. We observed that haemolysed/post-op plasma was red in colour due to the presence of fHb. Similarly, fHb-A_0_ solutions showed increasing intensity of colour with increasing fHb-A_0_ concentration, supported by cell lysates following treatment removal shown in Figure 2C. These results suggested that the fHb in the samples may have affected the bioluminescent signal output of the viability assay via optical interference as has been observed with other types of assays [19-21].

**Figure 2.**
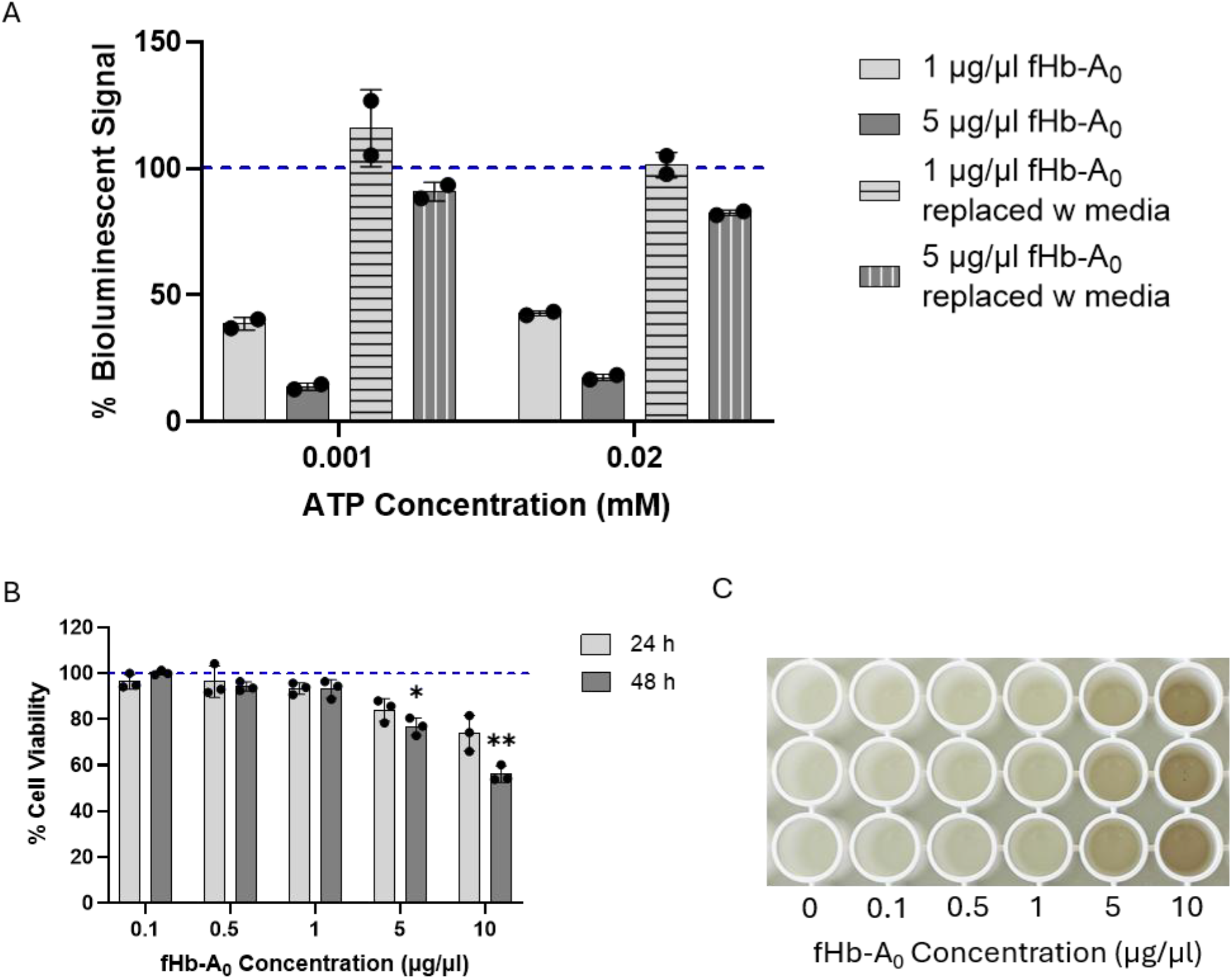
Investigating interference of free haemoglobin in a bioluminescent viability assay. (A) Solutions containing 1 and 5 µg/µl of free haemoglobin (fHb-A_0_) were incubated in a 96-well plate for 24 h. Following incubation, the fHb-A_0_ was removed from half of the wells and replaced with media. ATP was added to all wells, either at 0.02 or 0.001 mM, and the cell viability Cell Titer-Glo assay was used to quantify bioluminescent signal. Data presented is from 2 independent experiments, shown as mean % ± SD of the media control containing the respective ATP concentration. (B) ciPTECs were treated with fHb-A_0_ concentrations ranging from 0.1 to 10 µg/µl for 24 and 48 h. Treatment media was replaced with fresh media immediately prior to the use of the CellTiter-Glo assay to determine cell viability based on cellular ATP content. Data presented is from 3 independent experiments shown as mean % ± SD of the media control, which is shown by the dotted blue line. Statistical significance from the control at each time point was determined by One-way ANOVA followed by Dunnett’s multiple comparisons test and is represented by *, p<0.05, **, p<0.01. Controls are indicated by dotted blue lines. (C) Representative image of cell lysates after 24 h treatment with fHb-A_0_ following measurement of bioluminescent signal.

### fHb interfered with bioluminescent assays

To confirm that the presence of fHb itself interfered with the bioluminescent assay, we quantified ATP levels of solutions containing known concentrations of ATP (0.001 mM and 0.02 mM) after 24 h incubation with 1 and 5 µg/µl fHb-A_0_, in the absence of any cells. The results showed a reduction in bioluminescent signal output in the presence of fHb-A_0_, while wells in which the respective fHb-A_0_-treatment solution had been replaced with fresh media before the addition of ATP gave a much higher signal output. Specifically, the 1 µg/µl fHb-A_0_ solution reduced bioluminescent signals by 77.2 % and 58.6 % in the presence of 0.001 mM and 0.02 mM ATP, respectively (Figure 2A). Similarly, the 5 µg/µl fHb-A_0_ solution reduced bioluminescent signals by 77.0% and 64.8% in the presence of 0.001 mM and 0.02 mM ATP, respectively. Unlike the 1 µg/µl fHb-A_0_ solutions, even following the replacement of 5 µg/µl fHb-A_0_ with fresh media, the bioluminescent signal output did not return to control levels (Figure 2A). This was most likely due to fHb-A_0_ remaining in the wells which was then released into solution during incubation with the ATP lysis reagent and therefore reduced the bioluminescent signal measured.

Next, we determined whether replacement of fHb-A_0_-containing media prior to the viability assay mitigated fHb interference. We treated ciPTECs for 24 and 48 h with a range of fHb concentrations and replaced with fresh media immediately prior to the assay (Supplementary Figure S2), which resulted in concentration-dependent reductions in ciPTEC viability at both time points (Figure 2B). However, ciPTEC viability remained above 90% at concentrations from 0.1 to 1 µg/µl fHb-A_0_. Larger viability reductions were seen following 5 and 10 µg/µl fHb-A_0_ treatment which were significant at 48 h (Figure 2B). However, despite the replacement of fHb-A_0_ treatment solutions with fresh media prior to cell lysis, lysates following 24 h treatment with 5 and 10 µg/µl fHb-A_0_ remained visibly red (Figure 2C), suggesting that the fHb-A_0_ could not be sufficiently washed off with the media change. Thus, while we could establish a modest link between cell viability and fHb concentrations when fHb-A_0_ solutions had been replaced before performing the viability assay (Figure 2B), the reductions in cell viability were much smaller than without replacement of fHb-A_0_ solutions (Figure 1A). Together, these data indicated that the presence of fHb-A_0_ reduced the bioluminescent signal output via optical interference, and that fHb-A_0_ did not have the major effect on ciPTEC viability as initially indicated.

### Haemolysed plasma increased ciPTEC viability and stimulated NF-ĸB signalling

To re-evaluate the effect of haemolysis on ciPTEC viability, we treated the cells for 24 and 48 h with manually haemolysed plasma from healthy volunteers (different cohort to that in Figure 1). Besides baseline and haemolysed plasma, we also included 50:50 (baseline:haemolysed) mixed plasma for all volunteers, to create a gradient of fHb concentrations. Unlike after treatment with the fHb-A_0_ solutions, lysates from plasma-treated cells were not discoloured and this was reflected in the results. Interestingly, we observed a concentration-dependent increase in ciPTEC viability after both treatment periods. However, we detected lower viability when ciPTECs had been incubated with plasma for 48 h, independent of haemolysis, compared to 24 h. While baseline plasma resulted in a viability of 101.9 % ± 1.4, mixed and haemolysed plasma increased viability to 109.3 % ± 1.3 and 112.5 % ± 3.3, respectively at 24 h but this was not statistically significant (Figure 3A). Similarly, at 48 h mixed and haemolysed plasma significantly increased ciPTEC viability to 98.0 % ± 1.6 and 101.5 % ± 1.0, respectively, from 91.0 % ± 2.8 with baseline plasma (Figure 3B). Therefore, overall, haemolysed plasma did not reduce but rather increased cell viability, especially after 24 h treatment.

**Figure 3.**
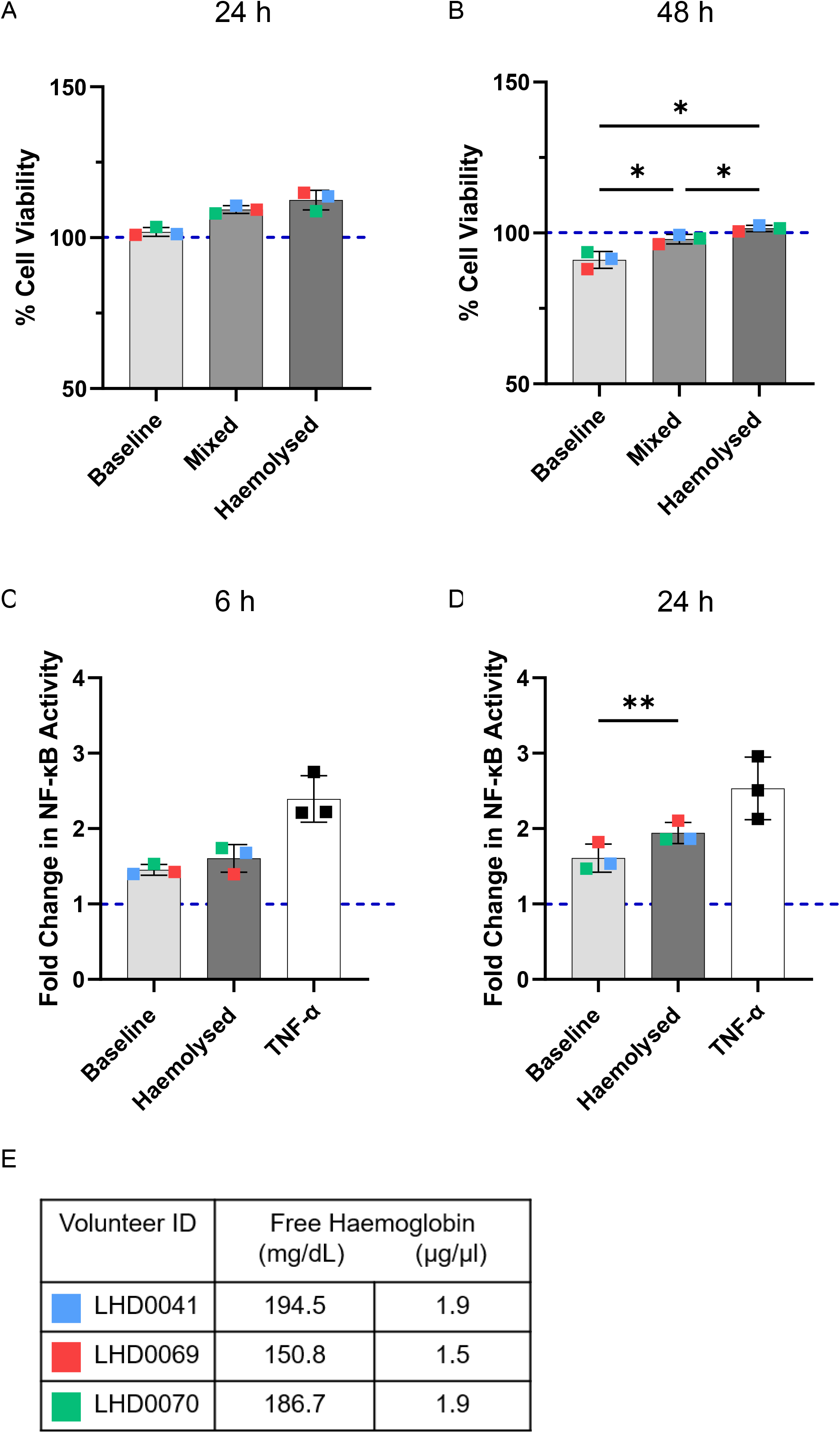
Comparing the effects of baseline and haemolysed healthy volunteer plasma on ciPTEC viability and NF-ĸB activity. ciPTECs were treated with baseline, mixed (a 1:1 mixture of baseline and hemolysed plasma) and haemolysed plasma from the same 3 healthy volunteers for (A) 24 h and (B) 48 h. The CellTiter-Glo assay was used to determine cell viability based on cellular ATP content. Statistical significance was determined by One-way ANOVA followed by Tukey’s multiple comparisons test and is represented by *, p<0.05. NF-ĸB ciPTECs were treated with baseline or haemolysed plasma from the same healthy volunteers and 100 ng/ml TNF-α for (C) 6 h and (D) 24 h. NF-ĸB activity was quantified by luciferase assay. Statistical significance was determined by paired t-test and is represented by **, p<0.01. All data presented is from 3 independent experiments shown as mean % ± SD of the media control, represented by the dotted blue line. Individual data points represent individual volunteers and are colour coded. (E) Free haemoglobin concentrations of the haemolysed volunteer plasma samples used in this figure measured by Hemcheck.

To provide an independent read-out for any effect of fHb-induced injury in the ciPTECs, we measured NF-ĸB activity using a ciPTEC-NF-ĸB reporter line. In this assay, luciferase activity was used as a read-out of NF-ĸB activity in response to haemolysed plasma. After 24 h treatment, haemolysed plasma induced a small but statistically significant increase in NF-ĸB activity compared to baseline plasma (haemolysed plasma: 1.9 ± 0.1-fold change to media control; baseline plasma: 1.6 ± 0.2-fold change, Figure 3D). Although statistically significant, the biological relevance of this increase is unclear since TNF-α, a known activator of the NF-ĸB signalling pathway, led to a more than 2-fold (6 h) or 2.5-fold (24 h) increase in luciferase activity.

It is also important to note that for both cell viability and NF-ĸB activity, the highest concentration of fHb in the mixed or haemolysed plasma did not necessarily correlate to the largest increase in read-out, although values were not dissimilar (Figure 3E). This suggests that other confounding factors within the plasma could influence the cell response.

### Haemolysed plasma did not cause mitochondrial dysfunction in ciPTECs

To evaluate the effect of haemolysed plasma on mitochondrial function in the ciPTECs, we used the Seahorse XF Mito Stress Test. This also provided an alternative quantification of cellular ATP, specifically the rate of ATP-linked production, which did not rely on bioluminescence and therefore should not have been impacted by fHb interference.

Results showed that haemolysed plasma from only one volunteer, LHD0070, influenced mitochondrial function by causing a reduction in basal respiration and ATP production compared to the respective baseline plasma but these were not statistically significant (Figure 4A & C). However, mixed plasma from the same volunteer had no observable effect. Haemolysed plasma from all volunteers reduced spare respiratory capacity in the ciPTECs, which was significant between mixed and haemolysed plasma for volunteers LHD0041 and LHD0070 (Figure 4B). Similarly, haemolysed plasma from all volunteers reduced the proton leak, although this was not statistically significant (Figure 4D). Furthermore, the coupling efficiency was slightly increased by haemolysed plasma from all volunteers, however, significant increases were only observed between baseline and mixed plasma for samples from volunteers LHD0060 and LHD0041 (Figure 4E). These results indicate that haemolysed plasma may have minimally reduced the capacity of mitochondria to deal with stress. However, our analysis could not demonstrate that haemolysis resulted in overall mitochondrial dysfunction.

**Figure 4.**
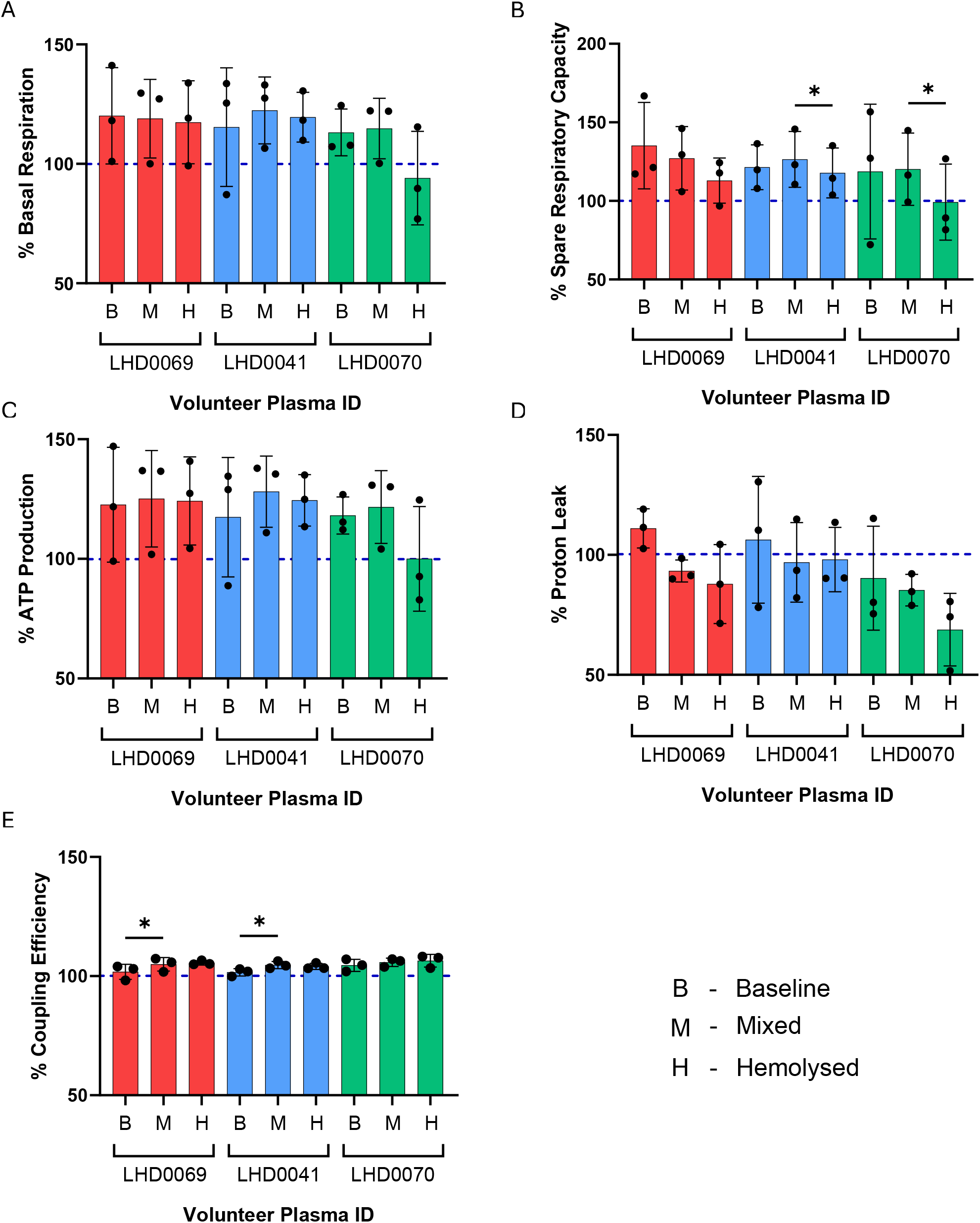
The effects of haemolysed volunteer plasma on ciPTEC mitochondrial respiration. The seahorse XF Mito Stress Test was performed on ciPTECs treated with baseline, mixed (a 1:1 mixture of baseline and manually haemolysed plasma) and manually haemolysed plasma from the same 3 healthy volunteers for 24 h. Analysis included (A) basal respiration, (B) spare respiratory capacity, (C) ATP production, (D) proton leak and (E) coupling efficiency. Data presented is from 3 independent experiments shown as mean % ± SD of the media control, indicated by the blue dotted line. Statistical significance between samples from the same volunteer were determined by One-way ANOVA followed by Tukey’s multiple comparisons test and are indicated by *, p<0.05.

## Discussion

In this study, we demonstrated that fHb interferes with bioluminescent signal output and therefore can compromise results from bioluminescent assays. Initial experiments suggested marked reductions in ciPTEC viability following exposure to fHb-A_0_ and haemolysed plasma. However, further investigation demonstrated that these effects were largely attributable to fHb-mediated assay interference. Following removal of this confounding effect, haemolysed plasma induced only modest changes in ciPTEC viability, inflammatory signalling and mitochondrial respiration. These findings highlight an important methodological consideration for studies investigating haemolysis-induced kidney injury.

Our analysis demonstrated that both fHb-A_0_ solutions and haemolysed plasma reduced bioluminescent signal output in a concentration-dependent manner. The viability assay relies on a firefly luciferase-generated bioluminescent signal used to quantify ATP content. The peak wavelength of bioluminescence emitted from the firefly luciferase enzymatic reaction at neutral pH is approximately 560 nm [22]. At this wavelength, Hb has a high absorbance and therefore may hinder the emittance of the true bioluminescent signal, causing optical interference [14]. A previous study found that Hb interfered with the emittance of luciferase luminescence in the context of tissue samples containing whole blood [23]. Removal of Hb from tissue samples resulted in a significant increase in the luminescent signal output. In the current study, in the absence of cells, the replacement of fHb-A_0_ solutions had concentration-dependent outcomes. The removal of treatment solutions containing low fHb-A_0_ concentrations was effective in restoring bioluminescent signal. However, this was not the case when replacing solutions containing high fHb-A_0_ concentrations as the bioluminescent signal was not fully recovered, possibly due to fHb adhering to the cell culture dish, and in the presence of cells, attaching to the cell surface or remaining within the cells [24, 25]. Interestingly, we did not observe fHb adherence when using haemolysed plasma despite observing an effect related to medium replacement. Together, these findings demonstrate that fHb can substantially reduce measured bioluminescent signals independent of cellular viability, leading to skewed results which significantly affect the interpretation of the fHb effect in experimental models of haemolysis.

Previous in vitro studies reported reduced viability following exposure of kidney cells to fHb solutions [26, 27]. The variations in the effect of fHb could be due to differences in cell models, their uptake of fHb, viability quantification methods and Hb interference. While our findings suggest that the ciPTECs show an injury response when exposed to fHb, overall our study indicates that the injury effect of fHb in kidney proximal tubule cells is rather mild. Therefore, it is possible that the magnitude of the effect of fHb solutions may have been overestimated when ATP-dependent bioluminescent assays were used without accounting for fHb interference.

In contrast to experiments using purified fHb-A_0_ solutions, treatment with haemolysed plasma from healthy volunteers was associated with small increases in ciPTEC viability and NF-κB activity. Human plasma naturally contains varying quantities of haptoglobin and hemopexin and it is therefore likely that some of the fHb and haem within the plasma following manual haemolysis is bound by these scavengers, limiting their effect on the cells [28, 29]. Consequently, total extracellular Hb concentrations measured photometrically using Hemcheck may not accurately reflect concentrations of biologically available unbound fHb [30]. Despite having a minimal effect on viability, haemolysed plasma induced evidence of cellular responses in line with previous reports of haemolysis-associated injury. Increased NF-ĸB activity is consistent with previous studies, in vitro and in vivo, which have shown that fHb or haem increase NF-ĸB activity at the protein or RNA level [31-33]. Similarly, we found that haemolysed plasma reduced mitochondrial spare respiratory capacity, suggesting reduced mitochondrial ability to adapt to further stress. However, in contrast to previous studies reporting more extensive mitochondrial dysfunction we did not observe evidence of marked impairment of mitochondrial respiration [34-37]. This may reflect the presence of endogenous scavenging proteins and antioxidants naturally found within human plasma that could mitigate the effects of fHb on the mitochondria [38].

There are a few limitations to this study. Firstly, as the replacement of solutions with high fHb concentrations did not restore full bioluminescent signal, it is not possible to determine to what extent the reductions in cell viability at these concentrations are due to cellular injury or remaining fHb. Secondly, the results of haemolysed plasma are based on data from only 3 healthy volunteers providing a very small sample size which all contain similar concentrations of fHb. Further investigation is required to validate our findings with a larger number of haemolysed plasma samples with a wide range of fHb concentrations. Thirdly, due to the photometric measurement of fHb concentration by the Hemcheck the unbound fHb concentrations in the plasma samples available to cause injury may not be accurate. It would be useful to compare Hemcheck concentrations with an assay which specifically binds unbound fHb to determine if the Hemcheck fHb concentrations are accurate. Regardless, the fHb concentrations measured in the manually haemolysed plasma likely exceed the binding capacity of the haptoglobin present in the plasma. Haptoglobin concentrations in human plasma range between approximately 30 and 200 mg/dL; we would therefore still expect to have seen an injury response [39, 40].

In conclusion, our study demonstrated fHb interference in a bioluminescent assay leading to initial overestimation of cytotoxicity. When this was accounted for, we demonstrated minimal impact on ciPTEC viability, NF-ĸB signalling and mitochondrial function in response to haemolysed plasma. These findings demonstrate the need for careful assay selection and mitigation of potential fHb interference in future investigations of haemolysis-induced kidney injury.

## Acknowledgments

We thank Professor Darius Widera, University of Reading, for providing the NF-ĸB vector plasmid, Dr Arthur Taylor for generating the viral particles used for the transduction and Professor Mark Field, University Hospitals Liverpool, for the use of clinical samples.

We wish to thank all patients and volunteers who agreed to take part in this study.

## Funding

Funding for this study was provided by the Alder Hey Children’s Kidney Fund to B.W., S.M., P.A.M., A.C., and by Kidney Research UK – Stoneygate to B.W., A.C., S.M., P.A.M.

## Conflicts of Interest

The authors declare no conflict of interest.

## Author Contributions

C.T., J.B., P.A.M., A.C., S.M., and B.W. conceived and designed research; C.T. performed experiments; C.T. analysed data; C.T. interpreted data; C.T. prepared figures; C.T. drafted manuscript; C.T., J.B., A.C., S.M., and B.W. edited and revised manuscript; C.T., J.B., P.A.M., A.C., S.M., and B.W. approved final version of manuscript.

## Ethical Approval

For the use of blood samples from aortic surgery patients, the study with the title: ‘Utilising point-of-care technology for the identification of haemolysis-induced Acute Kidney Injury following open surgery on the descending thoracic and thoracoabdominal aorta’, sponsored by Liverpool Heart and Chest Hospital NHS Trust, received ethics approval by the Hampstead Research Ethics Committee, IRAS ID 313578, REC reference 22/PR/0933, approval date 28/07/2022. All patients gave written informed consent.

For the use of volunteer blood, the study with the title: ‘Pharmacology Biobank 2022 – Collection and Storage of Human Samples of Biomedical Research’, sponsored by the University of Liverpool, received ethics approval by Central University Research Ethics Committee D, REC reference 11499, approval date 07/12/2022. All volunteers gave written informed consent.

## Use of AI

In the preparation of this manuscript and its data, there was no use of AI tools.

## Notes

### Competing Interest Statement

The authors have declared no competing interest.

