## Supplementary Files for "Analysis of haemolysis-based injury in immortalised proximal tubule epithelial cells is impaired by haemoglobin interference with bioluminescence assays"

**Supplementary Material**

**Supplementary Figure S2. Viability assay design**

Schematic showing the methods used for CellTiter-Glo or CellTiter-Glo 2.0 assays to determine and reduce free haemoglobin interference. Created in BioRender. Trott, C. (2026) https://BioRender.com/by0ct7d

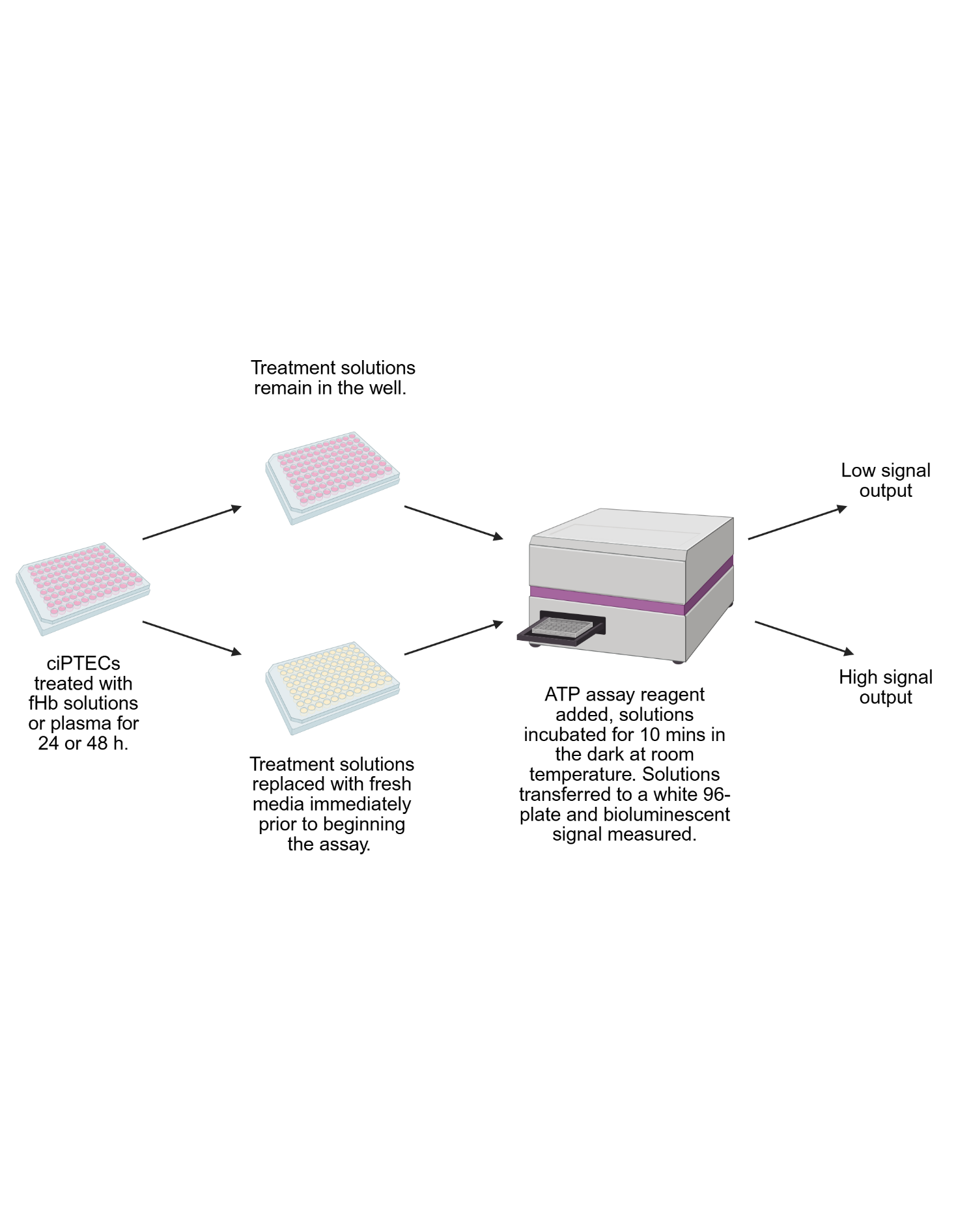

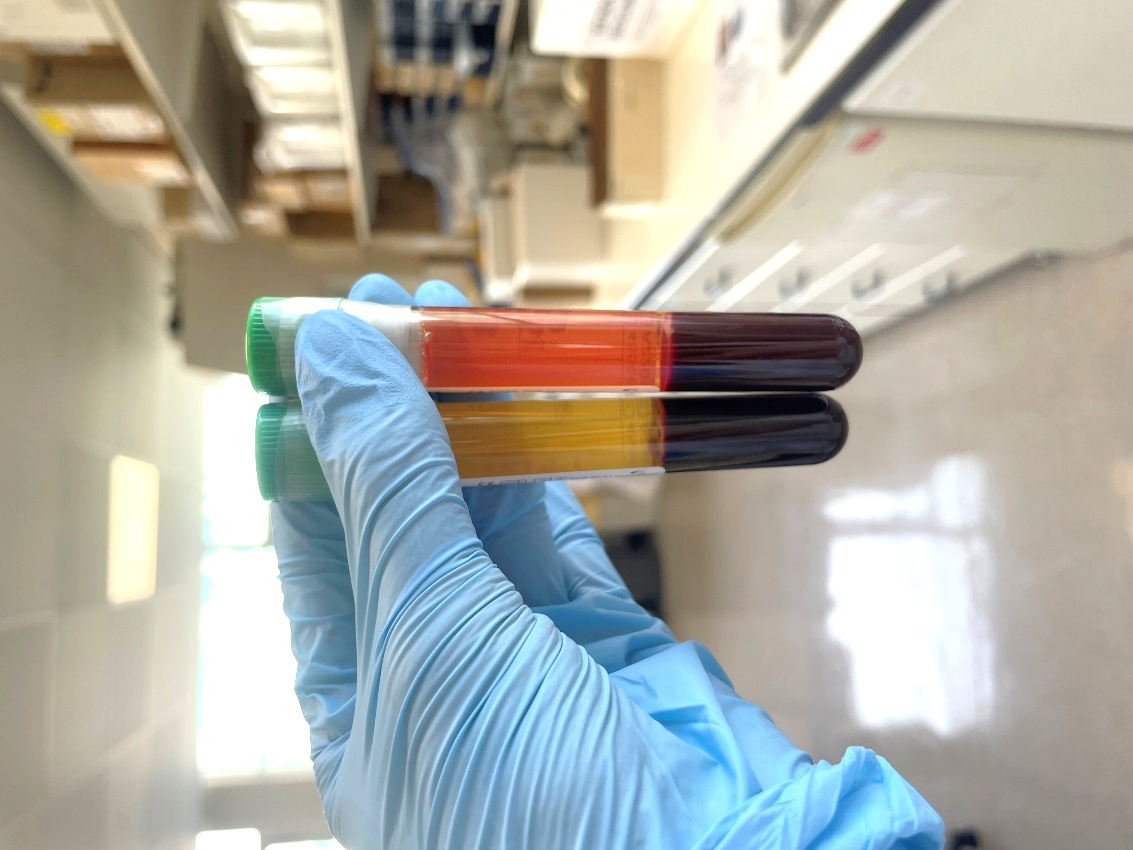

**Supplementary Figure S1. Volunteer blood sample following centrifugation.**

Image displaying volunteer blood samples following centrifugation. Image highlights the difference in plasma (the top layer), between manually haemolysed blood (left) and non-haemolysed blood (right) from the same volunteer.

**Supplementary Table S1.** Equations used for seahorse analysis.

| Parameter | Equation |
| --- | --- |
| Basal Respiration | (Last rate measurement before first injection) – (Non-mitochondrial respiration rate) |
| Spare Respiratory Capacity | (Maximal respiration) – (Basal respiration) |
| ATP Production | (Last rate measurement before oligomycin injection) – (Minimum rate measurement after oligomycin injection) |
| Proton Leak | (Minimum rate measurement after oligomycin injection) – (Non-mitochondrial respiration rate) |
| Coupling Efficiency | (ATP production rate) / (Basal respiration rate) x 100 |

**Supplementary Table S2.** Free haemoglobin concentrations measured by Hemcheck of patient post-op plasma samples shown in Figure 1D & E.

| Volunteer ID | Free Haemoglobin (mg/dL) (µg/µl) | |
| --- | --- | --- |
| HABHL0002 | 179 | 1.8 |
| HABHL0003 | 244 | 2.4 |
| HABHL0004 | 162 | 1.6 |
| HABHL0005 | 79 | 0.8 |
| HABHL0006 | 314 | 3.1 |
| HABHL0008 | 189 | 1.9 |
| HABHL0009 | 270 | 2.7 |
| HABHL0010 | 126 | 1.3 |
| HABHL0011 | 123 | 1.2 |
